# Ultra-deep sequencing reveals no evidence of oncogenic mutations or enrichment by *ex vivo* CRISPR/Cas9 genome editing in human hematopoietic stem and progenitor cells

**DOI:** 10.1101/2021.10.27.466166

**Authors:** M. Kyle Cromer, Valentin V. Barsan, Erich Jaeger, Mengchi Wang, Jessica P. Hampton, Feng Chen, Drew Kennedy, Irina Khrebtukova, Ana Granat, Tiffany Truong, Matthew H. Porteus

## Abstract

As CRISPR-based therapies enter the clinic, evaluation of the safety remains a critical and still active area of study. While whole genome sequencing is an unbiased method for identifying somatic mutations introduced by ex vivo culture and genome editing, this methodology is unable to attain sufficient read depth to detect extremely low frequency events that could result in clonal expansion. As a solution, we utilized an exon capture panel to facilitate ultra-deep sequencing of >500 tumor suppressors and oncogenes most frequently altered in human cancer. We used this panel to investigate whether transient delivery of high-fidelity Cas9 protein targeted to three different loci (using guide RNAs (gRNAs) corresponding to sites at AAVS1, *HBB*, and *ZFPM2*) at day 4 and day 10 timepoints post-editing resulted in the introduction or enrichment of oncogenic mutations. In three separate primary human HSPC donors, we identified a mean of 1,488 variants per Cas9 treatment (at <0.07% limit of detection). After filtering to remove germline and/or synonymous changes, a mean of 3.3 variants remained per condition, which were further reduced to six total mutations after removing variants in unedited treatments. Of these, four variants resided at the predicted off-target site in the myelodysplasia-associated *EZH2* gene that were subject to *ZFPM2* gRNA targeting in Donors 2 and 3 at day 4 and day 10 timepoints. While Donor 1 displayed on-target cleavage at *ZFPM2*, we found no off-target activity at *EZH2*. Sanger sequencing revealed a homozygous single nucleotide polymorphism (SNP) at position 14bp distal from the Cas9 protospacer adjacent motif in *EZH2* that eliminated any detectable off-target activity. We found no evidence of exonic off-target INDELs with either of the AAVS1 or HBB gRNAs. These findings indicate that clinically relevant delivery of high-fidelity Cas9 to primary HSPCs and *ex vivo* culture up to 10 days does not introduce or enrich for tumorigenic variants and that even a single SNP outside the seed region of the gRNA protospacer is sufficient to eliminate Cas9 off-target activity with this method of delivery into primary, repair competent human HSPCs.

## Main

The CRISPR system, consisting of a CRISPR/Cas protein coupled with a guide RNA (gRNA), has demonstrated remarkable versatility for site-specific genome editing. To ensure safe clinical translation of CRISPR systems for genome editing, insertions and deletions (indels) should occur only at the intended genomic site without off-target effects, through either non-homologous end joining (NHEJ) or homology-directed repair (HDR) pathways. Unintended genome editing occurs through low-fidelity Cas enzymes or when the gRNA directs cleavage to sequences similar to the target sequence, leading to the incorporation of off-target mutations that may have oncogenic or otherwise deleterious consequences.

Several recent reports have shown that DNA double-strand breaks (DSBs) introduced by Cas9 initiate a p53 response in pluripotent and cancer cell lines that results in cell cycle arrest and/or apoptosis^1,2^. Because cells with loss-of-function mutations in p53 do not suffer the same degree of toxicity following genome editing and may proceed through the cell cycle with unresolved DSBs, these studies suggested that Cas9-mediated cleavage can enrich for p53 mutations. However, the findings from these studies depended on: 1) the presence of p53 mutations in the initial pool of cells prior to (not as a consequence of) Cas9 delivery, which would not be expected to occur in primary cells derived from healthy donors; 2) stably integrated Cas9 expressed by a strong, constitutive promoter, which reliably invoke a dramatic DNA damage response^3^; and 3) immortalized cell lines that typically have gross chromosomal abnormalities (polyploidy, aneuploidy, translocations, etc.) with dysfunctional DNA damage and nucleic acid delivery-sensing responses^4–6^.

Significant efforts have thus been directed at not only predicting possible off-target genomic coordinates a priori^7–9^, but also at the development of empirical wet lab-based methods for identifying sites of off-target activity following genome editing^10–14^. While these experimental methods reported activity at many candidate sites, which may be missed by *in silico* prediction methods, there is some concern that these techniques depart from Cas9 delivery in a clinical setting since these methods typically involve long-term constitutive expression of wild-type Cas9 in immortalized cell lines or cell-free genomic DNA (gDNA). Therefore, there is an urgent need to assess the performance of these prediction algorithms and empirical methods in more therapeutically relevant contexts (i.e., via transient ribonucleoprotein (RNP)-based delivery of high-fidelity Cas9^15^ to human primary cells *ex vivo*).

The importance of long-term safety of genome editing/gene therapy in the clinic was illustrated recently when two sickle cell disease gene therapy trials (NCT02140554 and NCT04293185) were paused after two patients developed myeloid malignancies from either cytotoxic conditioning chemotherapy or insertional mutagenesis of the lentiviral vector^16^. Because of these safety concerns, in this study we sought to determine if oncogenic variants are introduced during Cas9 editing and/or the *ex vivo* expansion workflow. The ideal methodology necessitates targeted deep sequencing, however mutations with a variant allele frequency (VAF) below 1% remain mostly undetected by current genome-wide off-target detection techniques. This is in part because the signature of Cas9 nuclease activity is a spectrum of indels rather than primarily single nucleotide variants (SNVs). Therefore, an ultra-deep sequencing workflow capable of detecting SNVs as well as indels, amplifications, and multi-nucleotide variants (MNVs) has the potential to dramatically increase sensitivity for detection of the full spectrum of oncogenic off-target editing activity from 1% to <0.1% VAF, which will be necessary to identify low frequency variants that could promote pathogenic clonal expansion.

## Results

### Novel sequencing pipeline attains high coverage of tumor suppressors and oncogenes

To perform ultra-deep sequencing of tumor suppressors and oncogenes, we used a hybrid-capture next-generation sequencing (NGS) assay for detection of DNA variants at high depth across the exons of 523 cancer-relevant genes (spanning 1.94 Mb) using unique molecular indexes (UMIs) (named TruSight Oncology 500 (TSO500))^17^. These genes comprise known oncogenes in key guidelines of the most common cancer types, spanning non-small cell lung cancer to pancreatic adenocarcinoma (Supplemental Table 1). Prior work has shown a high degree of concordance (both positive and negative agreement) between the TSO500 panel and whole exome sequencing for measurement of mutation burden (nonsynonymous mutations per kilobase of DNA)^17^.

Hematopoietic stem and progenitor cells (HSPCs) from three separate healthy donors were subject to four conditions: Mock electroporated as well as three different Cas9 treatments with gRNAs targeting sites at AAVS1, *HBB*, or *ZFPM2* (Fig. 1A; Supplemental Table 2). Cas9 activity at AAVS1 and *HBB* have been thoroughly documented in the literature and these sites were chosen due to their relatively high and low off-target activity, respectively^11,12^, and the *HBB* gRNA is currently in phase I clinical trials for correction of the single SNP responsible for sickle cell disease^18,19^. As a positive control that we expected to elicit off-target activity in the TSO500 panel, we designed a gRNA targeting intron 3 of *ZFPM2*, which has a predicted off-target site in exon 5 of *EZH2*. This off-target site differs by a single nucleotide at position 1 of the protospacer, the site furthest from the PAM, which has the least bearing on Cas9 specificity (Fig. 1B), and is the highest ranked off-target site for the *ZFPM2* guide by COSMID^7^. *EZH2* was chosen as a relevant positive control because of its well-characterized role in a wide range of tumor types^20,21^ and the role of both loss- and gain-of-function mutations in myelodysplasias^22–25^, making it especially relevant for HSPC editing.

**Figure 1:**
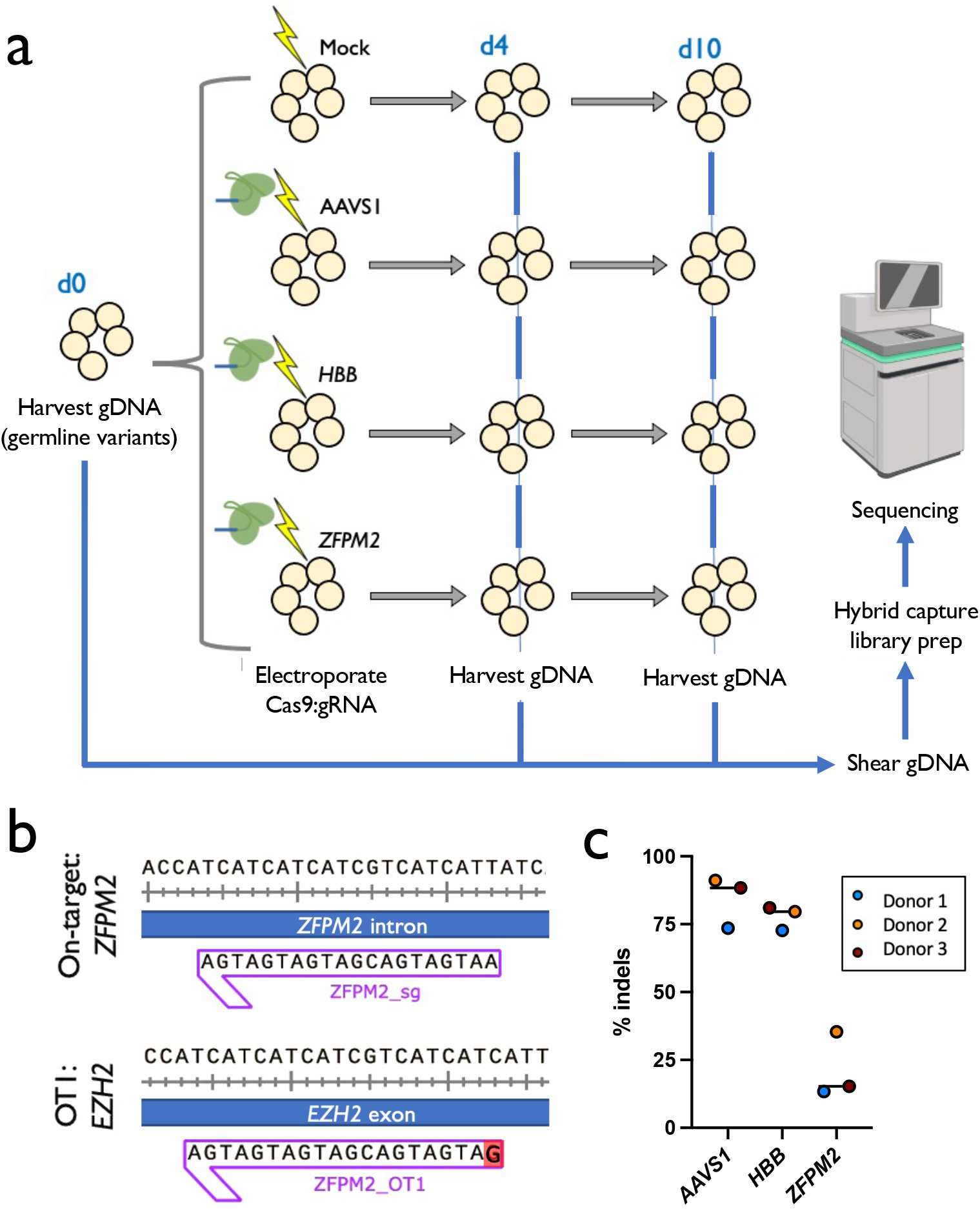
Experimental design & confirmation of on-target activity. a) Experimental design: CD34^+^ HSPCs from 3 donors underwent gDNA harvesting at d0 (to establish germline variants) and were then subject to mock electroporation or Cas9 treatments with gRNAs corresponding to sites at AAVS1, *HBB*, and *ZFPM2*. Cells were cultured and gDNA was harvested again at d4 and 10. b) Predicted off-target cut site (OT1) of *ZFPM2* guide in exon 5 of the *EZH2* gene, based on sequence homology. Mismatch in gRNA is shown in red. c) On-target activity of AAVS1, *HBB*, and *ZFPM2* gRNAs determined by PCR amplification of the genomic region surrounding the predicted cut sites followed by Sanger sequencing and analysis of indels by TIDE 4 days post-targeting. Bars represent median. N=3 separate HSPC donors.

The TSO500 genome editing workflow was adapted from a formalin-fixed paraffin-embedded (FFPE) tissue workflow to gDNA harvested from primary CD34^+^-purified umbilical cord blood-derived HSPCs from three separate healthy donors. Frozen cells were thawed and expanded for 2 days in HSPC media at 100K cells/mL and then targeted in the four treatment groups (2-5×10^5^ cells per treatment group) as reported previously^18,19,26,27^ (Fig. 1A). Genomic DNA was then harvested from 3-4×10^5^ cells at day 0 to establish germline variants and then cells were split into treatment groups, electroporated, and re-plated in fresh media. Because prior reports have shown that indel formation reaches completion 4 days after electroporating HSPCs with Cas9 RNP^27^, we harvested 4×10^5^ cells from each treatment group at day 4 and extracted gDNA for analysis. To determine whether enrichment of tumorigenic variants was occurring in our *ex vivo*-expanded HSPC populations, as well as to gain insight into whether *ex vivo* expansion itself (independent of Cas9 activity) was enriching for tumorigenic variants, we also harvested gDNA from the remainder of cells at 10 days post-targeting.

To ensure that high levels of on-target activity occurred for each gRNA, we performed targeted PCR amplification of the genomic region surrounding the predicted cut site followed by Sanger sequencing and analysis of indels by TIDE^28^. Indeed, a high frequency of on-target indels were observed across all three donors for AAVS1 and *HBB* gRNAs (Fig. 1C). While consistent across all donors, the *ZFPM2* gRNA induced fewer indels, which is not surprising due to its high degree of predicted off-target activity as well as the fact that this guide was not screened for efficiency prior to inclusion in this study, in contrast to the AAVS1 and HBB gRNAs that were identified as high activity gRNAs after screening multiple guides.

Pilot experiments were performed to confirm that the pipeline may be adapted from FFPE-derived tissue to gDNA harvested from primary cells. To determine the optimal amount of DNA for application to the sequencing pipeline, a range of 10-30ng of DNA was used as input for library preparation using the hybrid capture-based TSO500 Library Preparation Kit. Reads were mapped to the human genome (build hg19) and raw sequencing data was processed through a custom bioinformatic pipeline (Supplemental Fig. 1A) to identify indels, SNVs, and MNVs. Pilot experiments confirmed successful adaptation of the sequencing pipeline to gDNA harvested from primary cells in culture, and that at least 30ng of input DNA was necessary to achieve a median exon coverage (MEC) of 2000 (Supplemental Fig. 1B). To simultaneously detect intended edits, we supplemented the TSO500 panel with probes specific to AAVS1, *HBB*, and *ZFPM2* (Supplemental Table 3).

Following initial pilot experiments, raw sequencing data yielded a mean MEC >3550 for all samples per technical replicate, corresponding to a minimum limit of detection (LoD) and sensitivity of 0.205% and 95%, respectively (Fig. 2A; Supplemental Fig. 2). Moreover, because three technical replicates were sequenced for almost all timepoints and conditions, which are factored into the mean MEC of >3550, our LoD in these samples was further pushed to a limit of <0.07% VAF. Variants were consistent across technical replicates in terms of the types of variants called, with no significant differences comparing Mock to Cas9 treatments (Fig. 2B). We also observed a high degree of concordance across technical replicates, with a median of 98.31% of variants called in all replicates for each treatment for each donor (Fig. 2C; Supplemental Table 4). These data indicated that the total number of variants across replicates was more dependent on donor of origin than either time in culture or treatment with Cas9. In addition, the number of variants within each donor did not consistently increase due to time in culture (i.e., day 0 v. day 4 v. day 10) or treatment with Cas9. Consistent with these results, we found that read depth across the genome was more heavily influenced by donor than any other factor (Supplemental Fig. 3). In addition, while chromothripsis was recently reported as a rare consequence of on-target Cas9 cleavage^29^, in our bulk population of HSPCs we found no apparent drop in read depth in variants proximal to the intended cut site for any Cas9 treatment.

**Figure 2:**
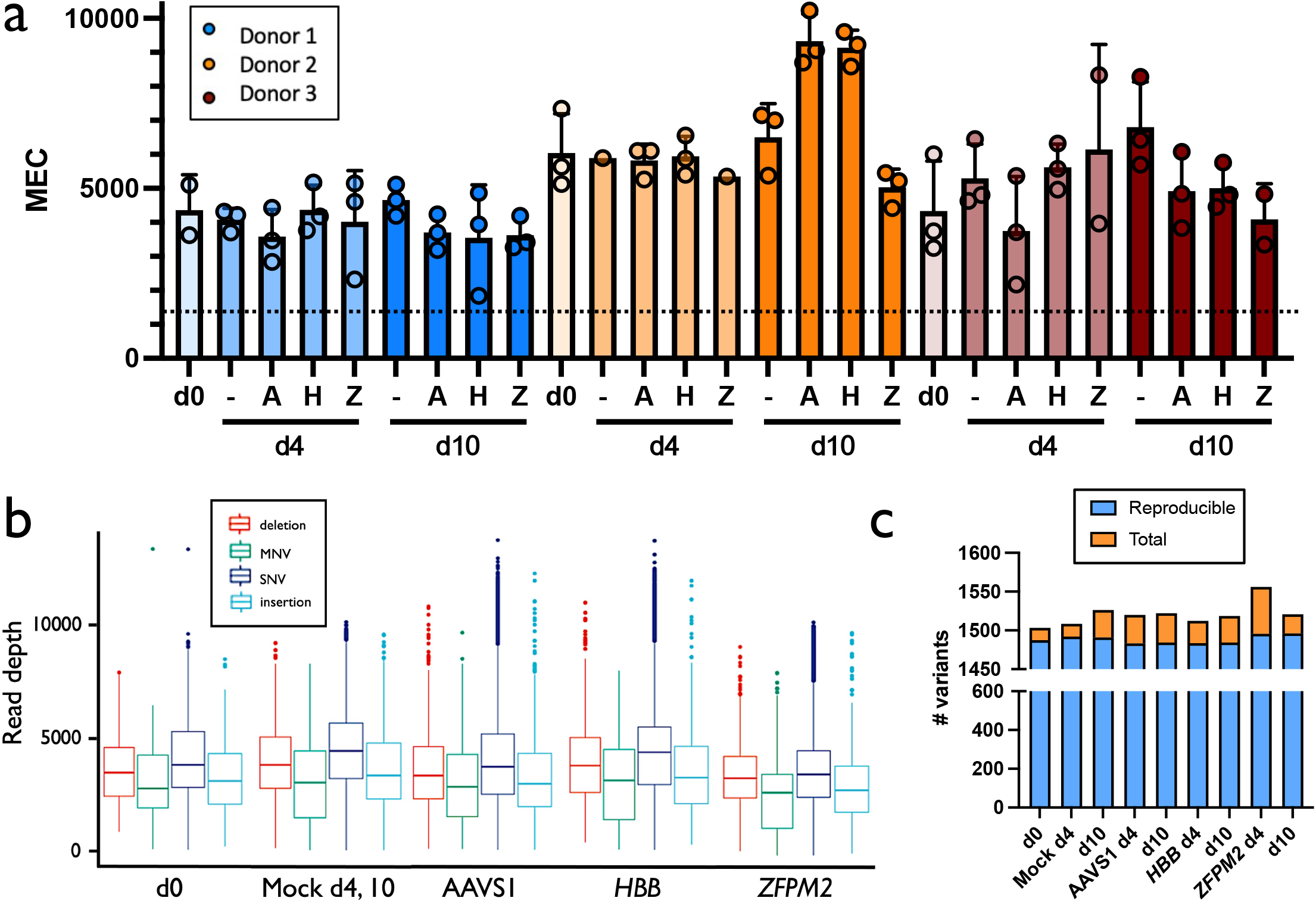
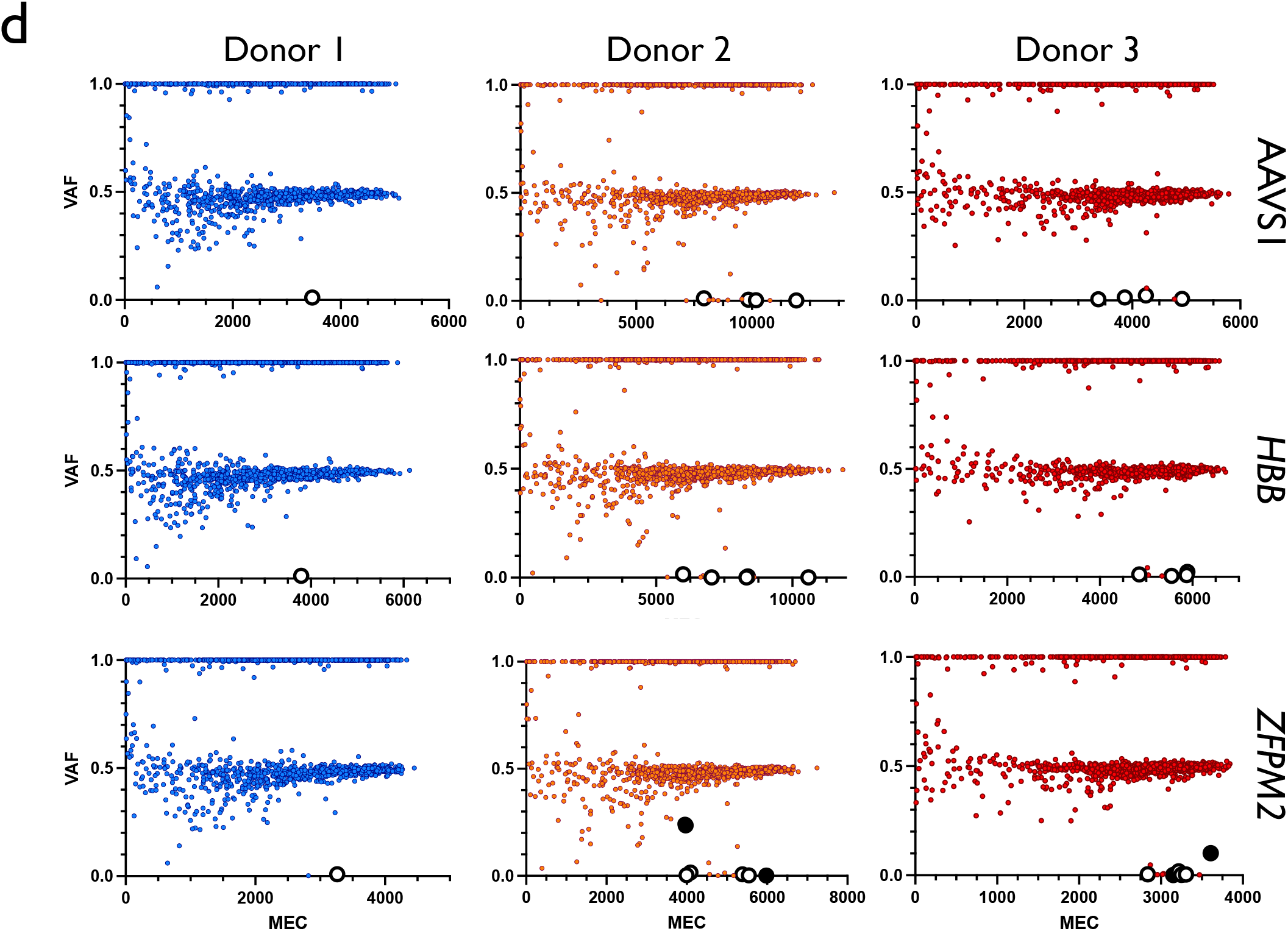
Summary of sequencing data. a) MEC for each treatment for each donor. Treatments are Mock electroporated (−), AAVS1-(A), *HBB-* (H), and *ZFPM2*-targeted (Z). Individual points represent technical replicates. Columns and error bars represent mean and standard deviation. b) MEC across all timepoints for each condition demonstrates consistent deep coverage for indels, SNVs, and MNVs. Midline represents median and box represents 25^th^ through 75^th^ percentiles. Variants among all donors and timepoints are combined for Cas9 treatments. In d0 and Mock, all donors are combined. c) Number of reproducible variants across technical replicates from total called by treatment group. Columns represent mean variants called for the three donors within each treatment. d) VAF x MEC for all variants found among technical replicates for Cas9 treatments for each donor at d10. Large white points are those that remained after removing germline and synonymous variants. Large black points are those that remain after removing variants present in Mock within each donor.

### Few variants found in treatments after filtering non-pathogenic germline mutations

To gain insight into the characteristics of the variants identified in our cohort, we plotted VAF by MEC for Mock samples at days 0, 4, and 10 (Supplemental Fig. 4A). Strikingly for all donors across all timepoints, the VAF frequencies trend toward 0.5 and 1.0 as MEC increases, which would correspond to heterozygous and homozygous germline variants, respectively. Because all variants were found within a panel of tumor suppressors and oncogenes, yet all HSPCs were derived from normal, healthy donors, we expect virtually all variants identified in Day 0 and Mock conditions to be non-pathogenic. Indeed, when filtering out both synonymous variants as well as those previously reported to occur >10 times in comprehensive germline databases^30,31^, an extremely small number of variants remain (a mean of 3.9 variants remaining from 1,490.1 reproducible variants per condition). Again, we found no consistent increase in the number of variants as HSPCs were cultured from d0 to d10. Interestingly, we observed several consistent variants that, while present in our germline database and consequently filtered, were found at intermediate VAFs rather than trending toward 0.5 or 1.0. While these variants were extremely consistent within, but not across donors, none of these were found in the exons of genes associated with clonal hematopoiesis of indeterminate potential^32,33^ (Note: the age of the donors for the source of the HSPCs is not known). Therefore, we believe these mutations represent either sequencing artefacts or bona fide HSPC donor chimerism that occurred prior to *ex vivo* culture or editing.

To determine whether editing with Cas9 introduces variants in tumor suppressors or oncogenes, we then plotted VAF x MEC for all Cas9 treatment groups at days 4 and 10 for all three donors (Fig. 2D; Supplemental Fig. 4B). Again, as expected for heterozygous and homozygous germline mutations, unfiltered variants trend toward VAFs of 0.5 and 1.0 as MEC increases. We next filtered out non-pathogenic variants by eliminating all called mutations that are synonymous and/or have been previously reported in the germline variant database. We found that our Cas9 treatments had a fewer number of variants remaining than even our Mock conditions (a mean of 3.3 variants remaining from 1,487.9 reproducible variants per condition). Because any variants also found in our Mock samples would not have been introduced by Cas9, we then removed all variants present in Mock conditions within each donor and only six variants remain among all eighteen Cas9 treatments (Fig. 3A). Of the six remaining variants, four of these are the expected *EZH2* mutations in Donors 2 and 3 within both day 4 and day 10 *ZFPM2* treatments. The other two variants that remain after filtering germline, synonymous, and Mock mutations are both SNVs found in d10 *ZFPM2* treatments in Donors 2 and 3 at <0.0015 VAF, which is close to our limit of detection. It is important to note that while only six variants remained in our treatment groups, the filters we applied to our Cas9 conditions were extremely conservative. Because Cas9 introduces indels far more frequently than SNVs at sites that display homology to the gRNA, if we apply these additional filters to our variants (i.e., remove SNVs as well as sites with no homology to the gRNA), only *EZH2* mutations remain.

**Figure 3:**
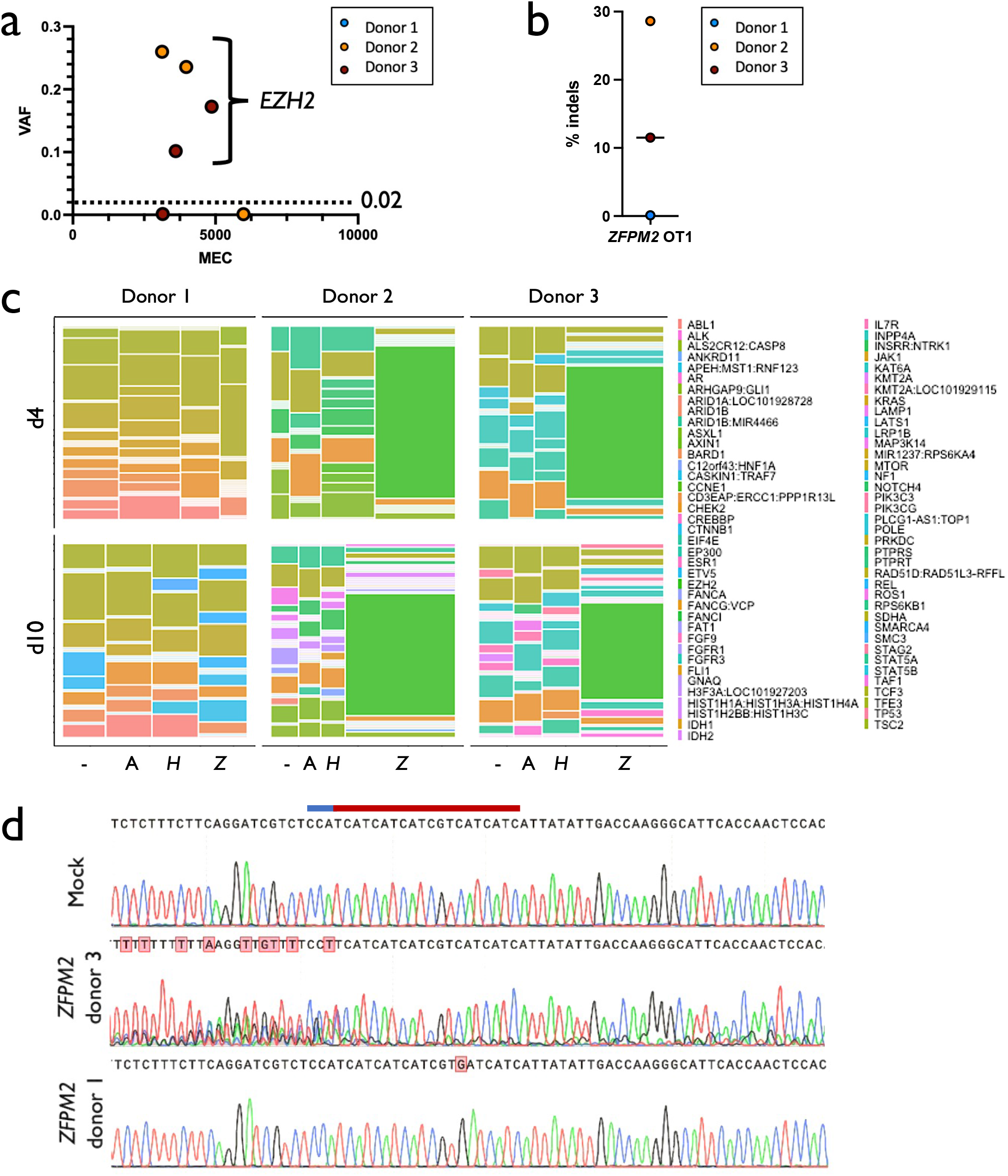
Variants identified in Cas9 treatments. a) VAF x MEC for variants remaining from all Cas9 treatments after removal of synonymous, germline, and Mock calls. Donor 1 had no variants remaining after filtering. b) Percent indels in *EZH2* identified by PCR amplification, Sanger sequencing, and TIDE analysis in d4 gDNA. Donor 1 had no detectable activity. c) Mosaic plot of genes harboring mutations within each donor and Cas9 treatment at d4 and 10. Area is proportional to the number of times variants were called in a particular gene within a particular treatment group. Filtering removed germline and synonymous variants. For each donor and timepoint, conditions are ordered as mock, AAVS1, *HBB*, and *ZFPM2* (−, A, *H*, and *Z*, respectively). d) Sanger chromatograms at predicted *EZH2* off-target site. The PAM site and protospacer are depicted as blue and red lines, respectively. Homozygous SNP in Donor 1 abrogated detectable editing activity.

### EZH2 off-target activity eliminated by homozygous SNP in Cas9 gRNA protospacer

In Donors 2 and 3, the expected *EZH2* off-target site displayed the highest VAF in both day 4 and day 10 timepoints across all three replicates at high confidence (3,893x average coverage) at an average of 19.3% off-target activity (Fig. 3A). Interestingly, the *EZH2* VAF in these donors decreased from day 4 to day 10 (mean of 21.7% to 16.9%, respectively), indicating a possible selective disadvantage for cells that harbor indels in this gene. The indel spectrum within *EZH2* was characterized (Supplemental Fig. 5) and total frequency was validated by PCR amplification, Sanger sequencing, and analysis of indels by TIDE (Fig. 3B). Notably, even without filtering Mock variants, mutations in *EZH2* comprised the vast majority of calls in Cas9 treatments in Donors 2 and 3 (Fig. 3C; Supplemental Fig. 6). Surprisingly, we found no detectable off-target activity at *EZH2* in Donor 1 by either NGS or TIDE (Fig. 3B) despite a high degree of on-target activity at *ZFPM2*. Upon investigation of the Sanger trace at this site in Donor 1, we found a homozygous SNP at position 6 of the protospacer. Due to the specificity of high-fidelity Cas9, which has been reported to reliably reduce off-target activity by 20-fold^15^, it is likely that this homozygous SNP eliminated all activity at this site (below the detection threshold of the TSO500 panel). The exceptional specificity of high-fidelity Cas9, when transiently delivered to primary cells, is evident from the single SNP outside of the core region of the protospacer in Donor 1 that was sufficient to eliminate all detectable activity at this site.

### Whole exome sequencing confirms absence of off-target activity from ex vivo culture & genome editing

Because transient delivery of Cas9 RNP and up to 10 days of ex vivo culture elicited few variants in the TSO500 panel, we next sought to expand our search for off-target activity to the entire exome. To do so, we electroporated high-fidelity Cas9 pre-complexed with AAVS1 gRNA to a single HSPC donor. We used the AAVS1 gRNA because it has been described as less specific than the HBB gRNA and we wanted to increase the chances of detecting any exonic off-target site. We then harvested gDNA from AAVS1-targeted and Mock electroporated treatments at d10 post-editing and subjected these to an exome capture panel and NGS. Prior to filtering, we identified 38,431 and 38,527 variants in Mock and AAVS1 treatments, respectively (Fig. 4A). To identify variants that may have resulted from Cas9 treatment, a tumor-normal pipeline was used to call somatic variants that were unique to the AAVS1 treatment (“tumor”) after subtracting the Mock as background (“normal”). In addition, we inverted the tumor-normal designation (i.e., treating Mock as tumor and AAVS1 treatment as normal) in order to estimate our background frequency of somatic calls resulting from this pipeline. These analyses identified 137 somatic variants in the AAVS1 treatment and 92 variants in the Mock condition (Fig. 4B). Because this pipeline is typically used to identify somatic variants in heterogeneous tumor samples, any mutation with a VAF notably greater than the “normal” sample was flagged. However, no off-target mutation introduced by Cas9 would have been present at any detectable VAF in the Mock condition. Therefore, we removed any variants found at >0.01 VAF in the Mock treatment, leaving 30 somatic mutations for further analysis (Fig. 4C). Though Cas9 nuclease activity typically introduces indels surrounding sites with a high degree of homology to the protospacer sequence, the majority of remaining variants after filtering (17/30) were SNVs and none of the 30 mutations were found to have >10bp match to the protospacer + PAM sequence within 20bp upstream or downstream of the called variant. We therefore conclude that neither targeted tumor suppressor/oncogene sequencing nor WES was able to identify any somatic mutations that occurred as a consequence of Cas9 activity. We therefore believe that the variants identified as somatic mutations in both Mock and AAVS1 treatments represent either real variation that occurred over the course of the 10 day ex vivo HSPC expansion or are sequencing artefacts.

**Figure 4:**
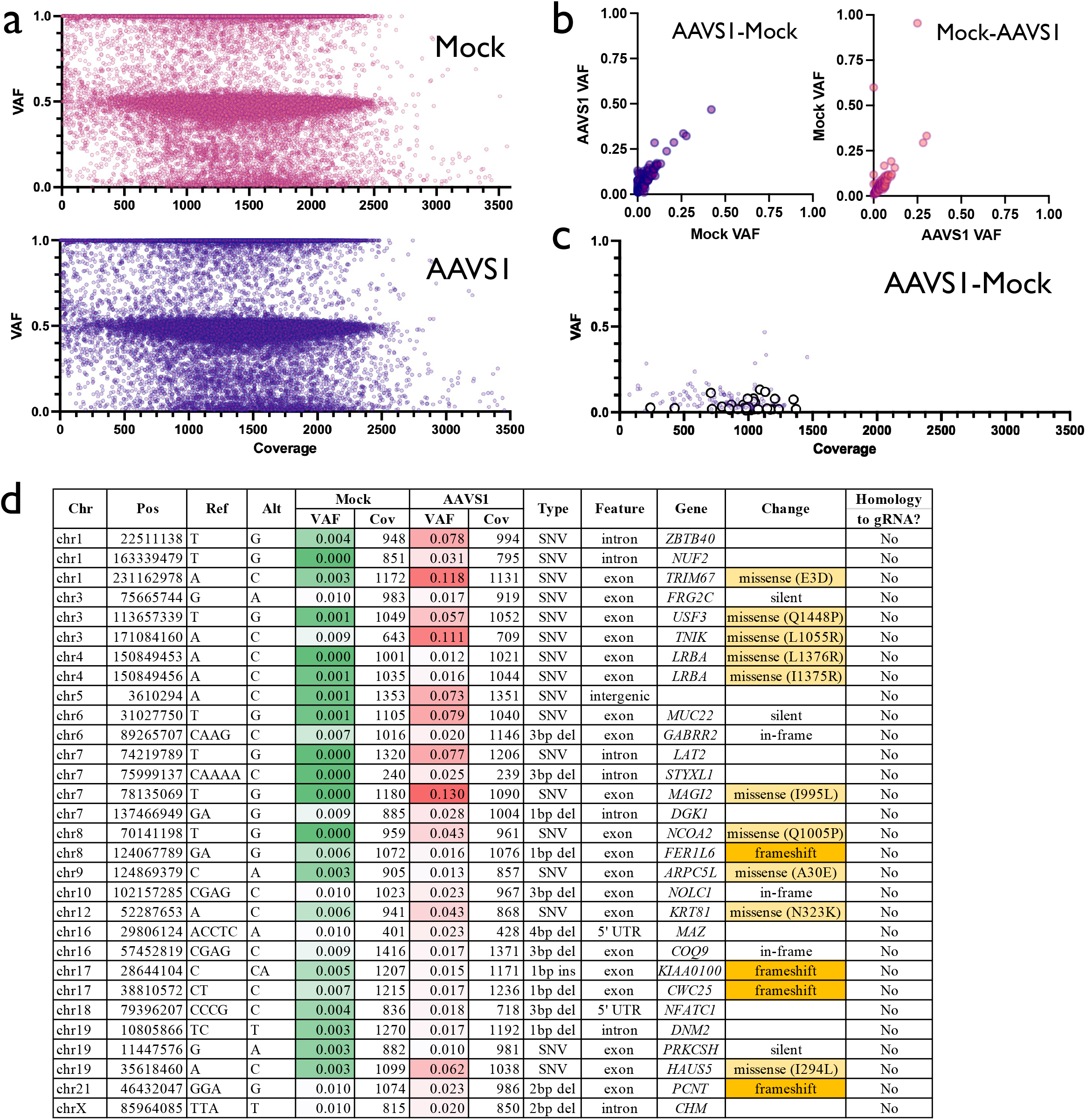
Variants identified in whole exome sequencing. a) VAF x Coverage for all variants called by exome sequencing pipeline in Mock and AAVS1 d10 conditions. Large white points are those that remained after removing germline and synonymous variants. Large black points are those that remain after removing variants present in Mock within each donor. b) VAF for variants called by tumor-normal pipeline when AAVS1 is used as tumor and Mock as normal inputs (left panel), and when Mock is used as tumor and AAVS1 as normal (right panel). c) VAF x Coverage for 137 variants shown in Panel B. 30 large white points are those that remained after removing variants with Mock VAF >0.01. d) Annotation for all 30 AAVS1 variants. Homology to gRNA is defined as 10 or more matches to protospacer+PAM within 20bp upstream or downstream of variant.

## Discussion

The first CRISPR-based therapies have entered early human clinical trials and many others are entering drug development pipelines^34^. There is a growing need to establish the long-term safety of edited human cells (*ex vivo* and *in vivo*) by CRISPR nucleases and vectors. Establishing the appropriate metrics for assessing genomic stability after genome editing continues to be an important and active area of study. For instance, whereas WGS is the only way to capture variants/abnormalities across the entire genome, read depth per base pair to achieve high sensitivity needed for genome editing purposes across a population of cells is not technically feasible. Sequencing the entire human genome only allows detection of high frequency events due to low per-base coverage. Alternatively, limiting sequencing to the most conserved and frequently mutated regions of the genome (e.g., exons, which comprises 1% of the genome^35^) allows for greater coverage and therefore greater detection power of lower frequency variants. Because cancer can occur due to expansion of even a single mutated clone, in this study we applied this concept to further limit sequencing to exons of the most common tumor suppressors and oncogenes to detect extremely low frequency events.

Cas9 is able to initiate DSBs in the DNA at both on- and off-target sites, potentially leading to unintended genomic abnormalities. In fact, several studies reported the enrichment of p53-inactivating mutations following CRISPR-based editing in immortalized human cell lines with constitutive or inducibly expressed Cas9 when a subset of p53 mutant cells were spiked in^1,2^. In contrast, prior studies in human primary cells found that Cas9 RNP delivery did not introduce mutations in p53 or 129 other cancer-related genes (using the Stanford Solid Tumor Actionable Mutation Panel)^36,37^. Therefore, we developed a novel tumor suppressor/oncogene ultra-deep sequencing pipeline to determine whether editing and short-term *ex vivo* expansion leads to disruption and/or enrichment of cancer-associated variants when delivered in a clinically relevant context—i.e., when high-fidelity Cas9^15^ is transiently delivered as RNP via electroporation to human primary HSPCs without subpopulations of cells with pre-existing tumorigenic variants. Toward this end, our workflow interrogated the exons of 523 known tumor suppressors and oncogenes and achieved levels of detection of germline and somatic mutations at <0.1% VAF.

When editing with three separate gRNAs (targeting AAVS1, *HBB*, and *ZFPM2)*, with ultra-deep sequencing of >500 tumor suppressors and oncogenes found no detectable variants (>0.002 VAF) that could be attributed to Cas9 activity or *ex vivo* expansion (aside from the expected *EZH2* off-target site in the *ZFPM2* treatment group). These findings were further confirmed by the absence of any off-target activity at sites resembling the AAVS1 gRNA by WES. In this clinically relevant context, transiently delivered high-fidelity Cas9 RNP into primary HSPCs did not introduce or enrich tumor variants. In fact, high-fidelity Cas9 was found to be so specific that even a single homozygous SNP at position 6 of the protospacer eliminated all detectable off-target activity in *EZH2*. In light of our findings, previous reports^1,2^ are likely an artefact of p53 mutant spike-in, genomic instability of cell lines, supraphysiological levels of Cas9 expression, and/or dramatic toxicity. Taken together, this work highlights the importance of: 1) regulating the duration and level of nuclease expression in order to limit the degree of off-target activity^12,38^; 2) minimizing toxicity through electroporation of RNP as opposed to mRNA- or plasmid-based editing^39^ so that opportunities for clonal expansion are minimized; and 3) conducting experiments in the most clinically relevant models—primary human cells with functional DNA sensing and damage repair machinery—rather than immortalized cell lines with well documented genomic abnormalities^4,5^.

A limitation of this work is that we focused entirely on the coding regions of genes, including those known to be involved in cancer. We chose this focus as off-target effects in coding regions, especially in tumor suppressor genes, might carry the highest risk for causing a serious adverse event. This focus, however, enabled high sequencing depth. Nonetheless, off-target INDELs in the non-coding region of the genome were not evaluated.

The importance of establishing safety of cell-based therapies prior to clinical translation is illustrated by the recent development of leukemia in two patients enrolled in a lentiviral gene therapy trial for sickle cell disease, which resulted in pausing of both related trials^16^. Follow-up investigation found that leukemic cells harbor viral integrations and that mutations in *RUNX1* and *PTPN11* occurred at some point during or following myeloablative conditioning and/or lentiviral integration. Disruption of both of genes have been shown to play a role in a wide variety of cancers^40–43^, and due to inclusion in the TSO500 panel and the sequencing depth we achieved in this study, we would have been able to identify variants in these genes at ≥0.1% VAF prior to autologous transplantation in these trials. Therefore, we believe that our study not only establishes an important benchmark for the typical degree of variation in cancer-associated genes following CRISPR-based editing and short-term *ex vivo* expansion, but also may become a common tool for assessing safety of cell-based products prior to transplantation (particularly in the event of clonal expansion and/or long-term *ex vivo* expansion). In doing so, we hope to maximize the longterm safety of the new generation of site-specific genome editing therapies. As CRISPR/Cas9 editing becomes more clinically widespread, identification and avoidance of genotoxicity will profoundly impact the pace with which these curative approaches reach patients safely.

## Acknowledgements

V.B. is an Anne T. and Robert M. Bass Endowed Fellow, supported by the Stanford Child Health Research Institute.

## Methods

### Culturing of HSPCs

Primary human HSPCs were sourced from fresh umbilical cord blood (generously provided by Binns Family program for Cord Blood Research) under protocol 33818, which was approved and renewed annually by the NHLBI IRB. All patients provided informed consent for the study. CD34^+^ HSPCs were bead-enriched using Human CD34 Microbead Kits (Mitenyi Biotec, Inc., Bergisch Gladbach, Germany) according to manufacturer’s protocol and cultured at 1×10^5^ cells/mL in CellGenix GMP SCGM serum-free base media (Sartorius CellGenix GmbH, Freiburg, Germany) supplemented with stem cell factor (SCF)(100ng/mL), thrombopoietin (TPO)(100ng/mL), FLT3–ligand (100ng/mL), IL-6 (100ng/mL), UM171 (35nM), 20mg/mL streptomycin, and 20U/mL penicillin. The cell incubator conditions were 37°C, 5% CO_2_, and 5% O_2_.

### Genome editing of HSPCs

Chemically modified sgRNAs used to edit HSPCs were purchased from Synthego (Menlo Park, CA, USA). The sgRNA modifications added were the 2’-O-methyl-3’-phosphorothioate at the three terminal nucleotides of the 5’ and 3’ ends described previously^38^. All Cas9 protein (SpyFi S.p. Cas9 nuclease) was purchased from Aldevron, LLC (Fargo, North Dakota, USA). The RNPs were complexed at a Cas9:sgRNA molar ratio of 1:2.5 at 25°C for 10min prior to electroporation. HSPCs were resuspended in P3 buffer (Lonza, Basel, Switzerland) with complexed RNPs and electroporated using the Lonza 4D Nucleofector (program DZ-100). Cells were plated at 1×10^5^ cells/mL following electroporation in the cytokine-supplemented media described above.

### TSO500 library preparation

Input DNA concentration was determined by Qubit dsDNA HS assay kit on the Qubit Fluorometer according to the manufacturing protocol (Qubit, London, UK). DNA was then fragmented to 90 to 250 bp by sonication using a Covaris E220 Evolution Sonicator (Covaris, Woburn, MA, USA), with a target peak of around 130bp as determined by Agilent Technologies 2100 Bioanalyzer using a High Sensitivity DNA chip. Samples then underwent end repair and A-tailing. Adapters containing UMIs were ligated to the ends of the DNA fragments. After a purification step, the DNA fragments were amplified using primers to add index sequences for sample multiplexing (required for cluster generation). Two hybridization/capture steps were performed. First, a pool of oligos specific to the 523 genes targeted by TSO500 with supplementary probes from IDT (Supplementary Table 3) were hybridized to the prepared DNA libraries overnight. Next, streptavidin magnetic beads were used to capture probes hybridized to the targeted regions. The hybridization and capture steps were repeated using the enriched DNA libraries to ensure high specificity for the captured regions. Primers were used to amplify the enriched libraries using sample purification beads. The enriched libraries were quantified and each library was normalized to ensure a uniform representation in the pooled libraries. Finally, the libraries were pooled, denatured, and diluted to the appropriate loading concentration and sequenced on an Illumina NovaSeq with a read length of 2×151 base pairs. Up to 8 TSO500 libraries were sequenced per run.

### Indel frequency analysis by TIDE

2-4d post-targeting, HSPCs were harvested and a Qiagen DNeasy Blood & Tissue Kit (Redwood City, CA, USA) was used to collect gDNA. The following primers were then used to amplify respective cut sites at with Phusion Green Hot Start II High-Fidelity PCR Master Mix (Thermo Fisher Scientific, Waltham, MA, USA) according to manufacturer’s instructions: AAVS1, forward: 5’-AGGATCCTCTCTGGCTCCAT-3’, reverse: 5’-CCCCTGTCATGGCATCTTC-3’; *HBB*, forward: 5’-AGGGTTGGCCAATCTACTCC-3’, reverse: 5’-AGTCAGTGCCTATCAGAAACCCAAGAG-3’; *ZFPM2*, forward: 5’-GCAAATGCAGCAGTAGACCA-3’, reverse: 5’-CCTTCGCTCTCAATTTTGCT-3’; and *EZH2 (ZFPM2* OT1), forward: 5’-AAAAGAGAAAGAAGAAACTAAGCCCTA-3’, reverse: 5’-TTTTCCTCCCCTCATTTCAA-3’. PCR reactions were then run on a 1% agarose gel and appropriate bands were cut and gel-extracted using a GeneJET Gel Extraction Kit (Thermo Fisher Scientific, Waltham, MA, USA) according to manufacturer’s instructions. Gel-extracted amplicons were then Sanger sequenced with the forward and reverse amplicon primers shown above. Resulting Sanger chromatograms were then used as input for indel frequency analysis by TIDE as previously described^26^.

### Whole-exome sequencing

Data was processed using the DRAGEN v3.8.4 Enrichment pipeline. All datasets were processed as independent samples, and related mock and test samples were additionally processed as “tumor/normal” and “normal/tumor” pairs. Known systematic noise filters were applied to all called variants.

---

**Supplemental Table 1:**
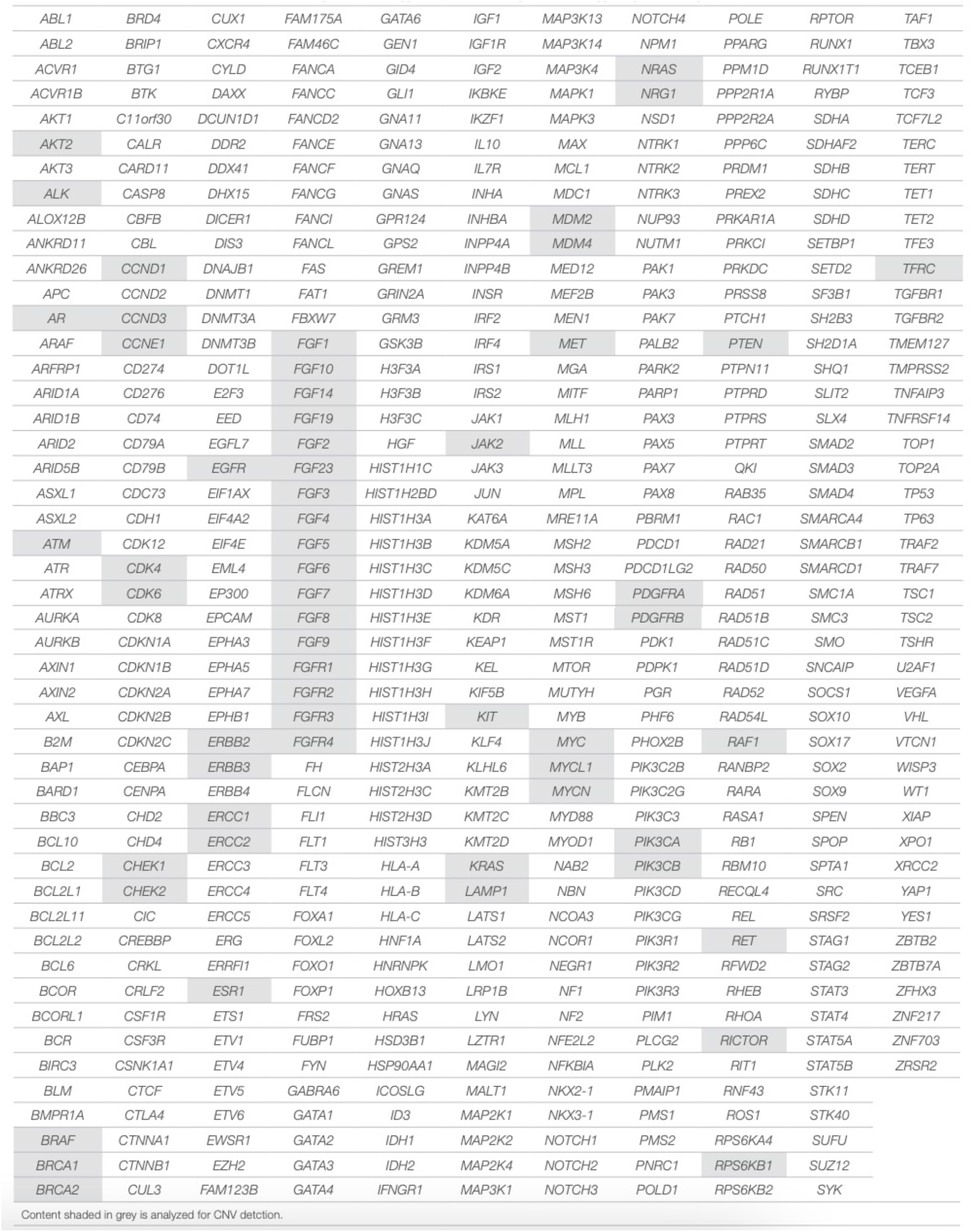
Genes included on TSO500 panel.

**Supplemental Table 2:**
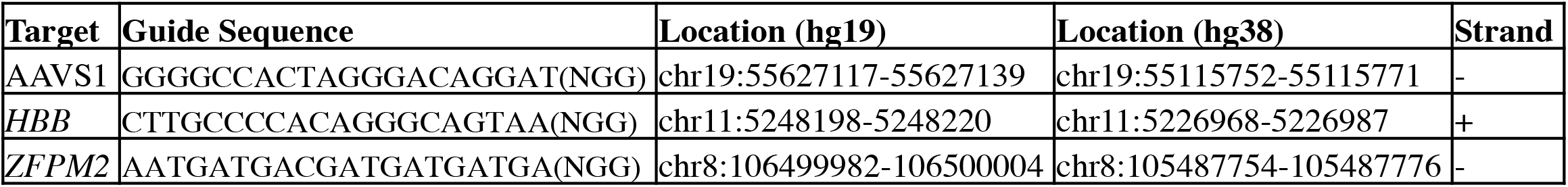
Cas9 gRNA information.

**Supplemental Figure 1:**
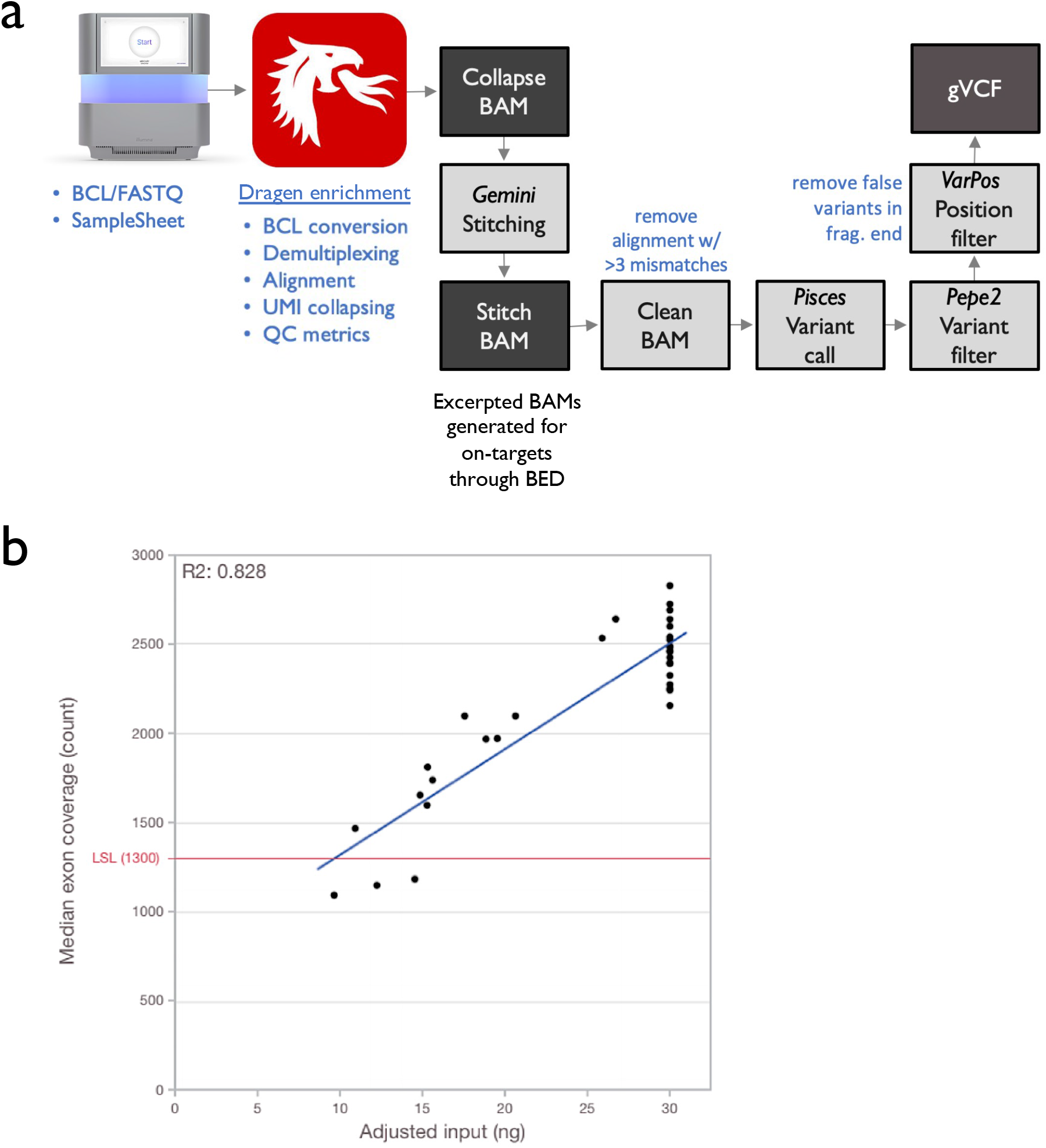
Bioinformatic pipeline used to align reads with genome. a) Bioinformatics pipeline used to map reads to genome, call, and annotate variants. b) DNA input and corresponding MEC yielded in pilot library prep and sequencing runs. LSL = recommended lower specification limit, set at 1300 MEC.

**Supplemental Table 3:** Supplementary probe information. 36 probes were designed for the two gene symbol inputs by IDT (*HBB* and *ZFPM2*). Bed coordinates are shown in hg19 assemblies. For gene capture panels (i.e., *HBB* and *ZFPM2*), the probes come from IDT’s xGen Exome Research Panel. They are spaced at 1x tiling where every base is covered by at least one probe. The original exome design was performed with repeat masking and empirically validated so we do not perform any additional QC to the probe sequences. For AAVS1, we padded probes 1kb upstream and downstream (120bp probes collectively hybridized to all exons with at least 1x tiling, meaning each base is covered by at least one probe). For SNPs, we padded out to 100bp with all SNPs directly in the middle. We then designed at 1x tiling without repeat masking and IDT performed predictive QC analysis to the probe sequences. The QC analysis involved standard BLAST and Minimap alignment.

| Chromosome | Start | Stop | Name |
| --- | --- | --- | --- |
| chr19 | 55626110 | 55626230 | AAVS1 |
| chr19 | 55626230 | 55626350 | AAVS1 |
| chr19 | 55626350 | 55626470 | AAVS1 |
| chr19 | 55626470 | 55626590 | AAVS1 |
| chr19 | 55626590 | 55626710 | AAVS1 |
| chr19 | 55626710 | 55626830 | AAVS1 |
| chr19 | 55626830 | 55626950 | AAVS1 |
| chr19 | 55626950 | 55627070 | AAVS1 |
| chr19 | 55627070 | 55627190 | AAVS1 |
| chr19 | 55627190 | 55627310 | AAVS1 |
| chr19 | 55627310 | 55627430 | AAVS1 |
| chr19 | 55627430 | 55627550 | AAVS1 |
| chr19 | 55627550 | 55627670 | AAVS1 |
| chr19 | 55627670 | 55627790 | AAVS1 |
| chr19 | 55627790 | 55627910 | AAVS1 |
| chr19 | 55627910 | 55628030 | AAVS1 |
| chr19 | 55628030 | 55628150 | AAVS1 |
| chr11 | 5246772 | 5246892 | HBB |
| chr11 | 5246892 | 5247012 | HBB |
| chr11 | 5247797 | 5247917 | HBB |
| chr11 | 5247917 | 5248037 | HBB |
| chr11 | 5248145 | 5248265 | HBB |
| chr8 | 106331145 | 106331265 | ZFPM2 |
| chr8 | 106431331 | 106431451 | ZFPM2 |
| chr8 | 106431451 | 106431571 | ZFPM2 |
| chr8 | 106456498 | 106456618 | ZFPM2 |
| chr8 | 106573590 | 106573710 | ZFPM2 |
| chr8 | 106646469 | 106646589 | ZFPM2 |
| chr8 | 106800929 | 106801049 | ZFPM2 |
| chr8 | 106801049 | 106801169 | ZFPM2 |
| chr8 | 106810943 | 106811063 | ZFPM2 |
| chr8 | 106811063 | 106811183 | ZFPM2 |
| chr8 | 106813260 | 106813380 | ZFPM2 |
| chr8 | 106813380 | 106813500 | ZFPM2 |
| chr8 | 106813500 | 106813620 | ZFPM2 |
| chr8 | 106813620 | 106813740 | ZFPM2 |
| chr8 | 106813740 | 106813860 | ZFPM2 |
| chr8 | 106813860 | 106813980 | ZFPM2 |
| chr8 | 106813980 | 106814100 | ZFPM2 |
| chr8 | 106814100 | 106814220 | ZFPM2 |
| chr8 | 106814220 | 106814340 | ZFPM2 |
| chr8 | 106814340 | 106814460 | ZFPM2 |
| chr8 | 106814460 | 106814580 | ZFPM2 |
| chr8 | 106814580 | 106814700 | ZFPM2 |
| chr8 | 106814700 | 106814820 | ZFPM2 |
| chr8 | 106814820 | 106814940 | ZFPM2 |
| chr8 | 106814940 | 106815060 | ZFPM2 |
| chr8 | 106815060 | 106815180 | ZFPM2 |
| chr8 | 106815180 | 106815300 | ZFPM2 |
| chr8 | 106815300 | 106815420 | ZFPM2 |
| chr8 | 106815420 | 106815540 | ZFPM2 |
| chr8 | 106815540 | 106815660 | ZFPM2 |
| chr8 | 106815660 | 106815780 | ZFPM2 |

**Supplemental Figure 2:**
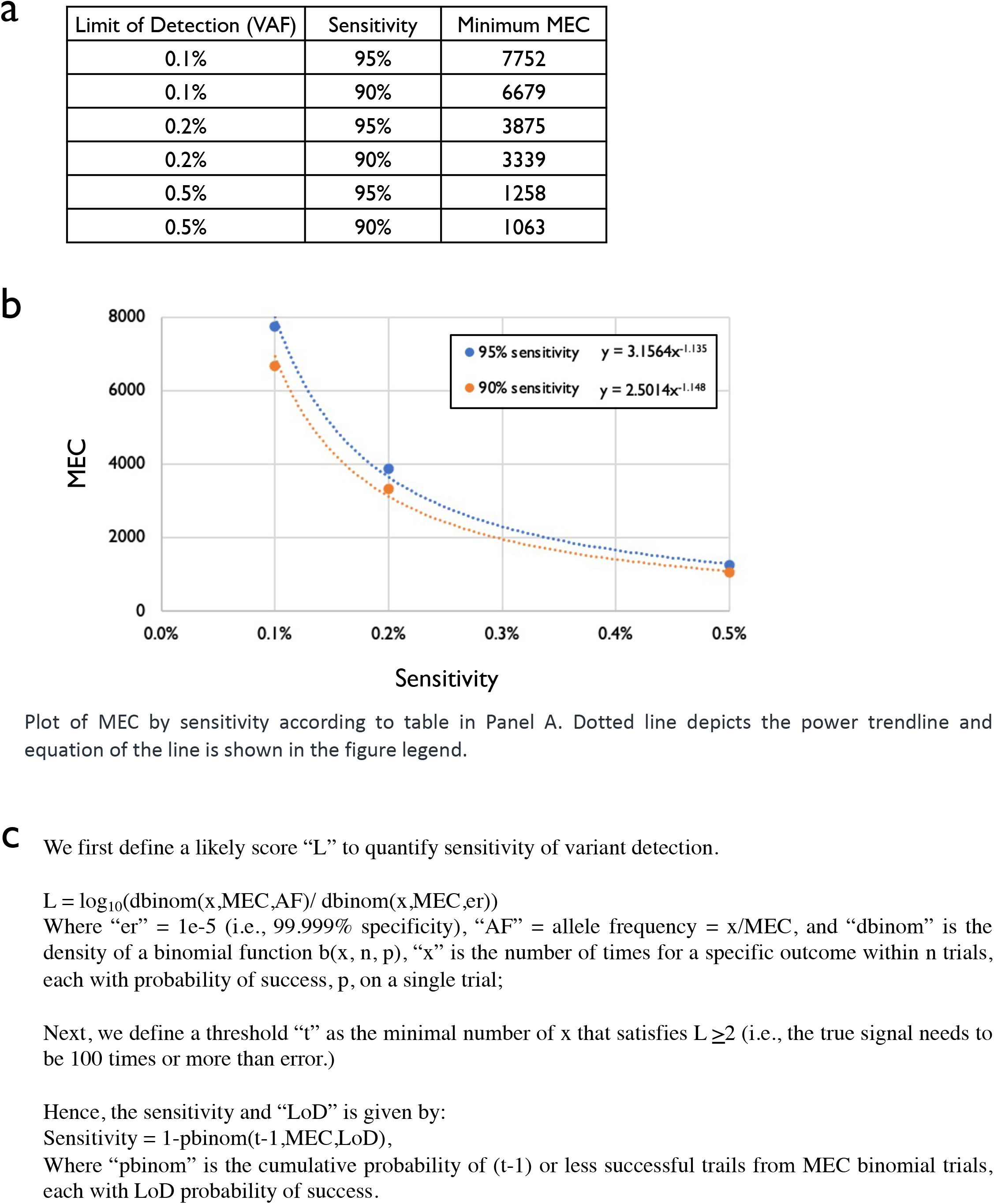
Calculation of limit of detection & sensitivity given MEC. Plot of MEC by sensitivity according to table in Panel A. Dotted line depicts the power trendline and equation of the line is shown in the figure legend. We first define a likely score “L” to quantify sensitivity of variant detection. L = log_10_(dbinom(x,MEC,AF)/ dbinom(x,MEC,er)) Where “er” = 1e-5 (i.e., 99.999% specificity), “AF” = allele frequency = x/MEC, and “dbinom” is the density of a binomial function b(x, n, p), “x” is the number of times for a specific outcome within n trials, each with probability of success, p, on a single trial; Next, we define a threshold “t” as the minimal number of x that satisfies L≥2 (i.e., the true signal needs to be 100 times or more than error.) Hence, the sensitivity and “LoD” is given by: Sensitivity = 1-pbinom(t-1,MEC,LoD), Where “pbinom” is the cumulative probability of (t-1) or less successful trails from MEC binomial trials, each with LoD probability of success.

**Supplemental Table 4:**
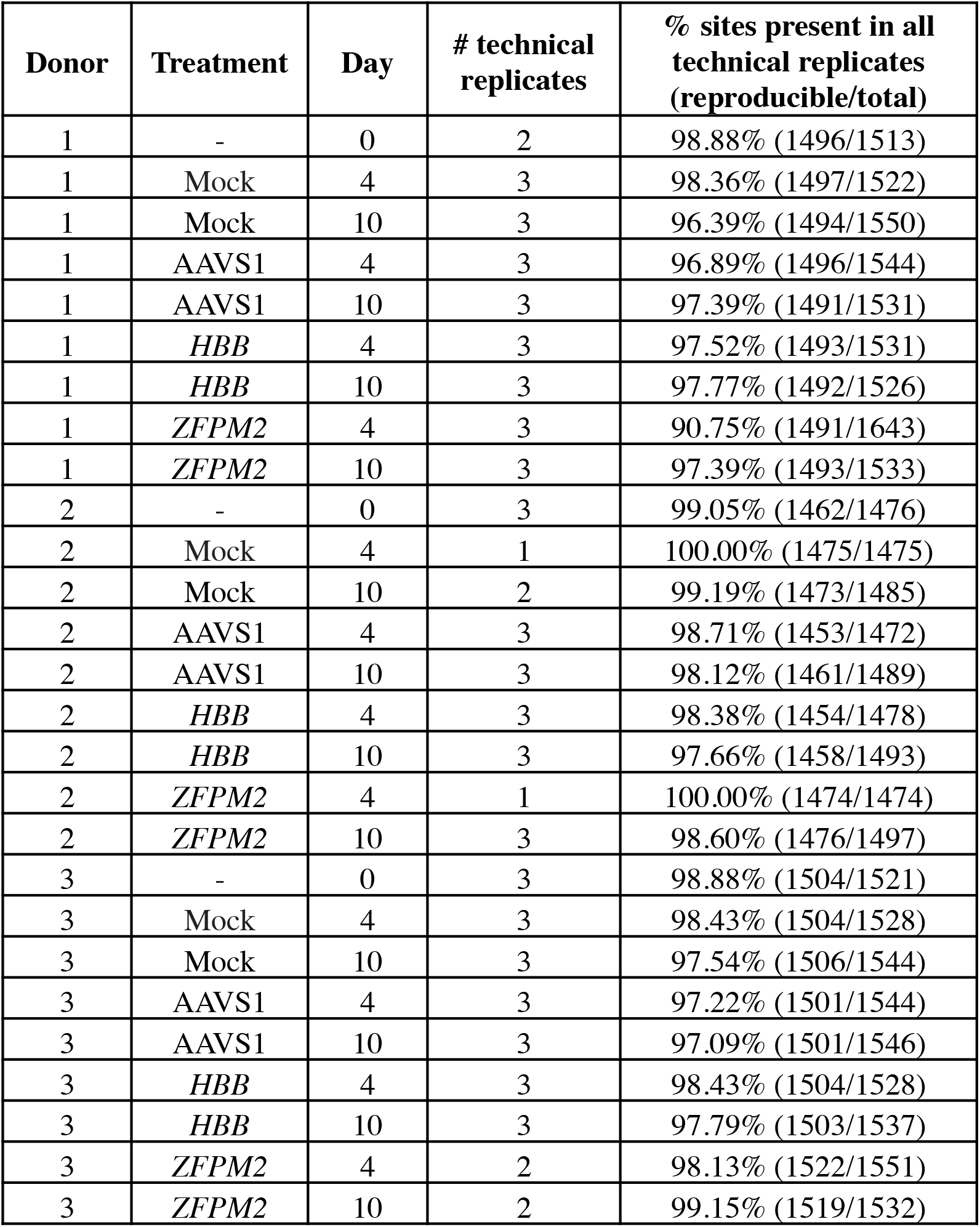
Reproducibility across technical replicates.

| Donor | Treatment | Day | # technical replicates | % sites present in all technical replicates (reproducible/total) |
| --- | --- | --- | --- | --- |
| 1 | - | 0 | 2 | 98.88% (1496/1513) |
| 1 | Mock | 4 | 3 | 98.36% (1497/1522) |
| 1 | Mock | 10 | 3 | 96.39% (1494/1550) |
| 1 | AAVS1 | 4 | 3 | 96.89% (1496/1544) |
| 1 | AAVS1 | 10 | 3 | 97.39% (1491/1531) |
| 1 | *HBB* | 4 | 3 | 97.52% (1493/1531) |
| 1 | *HBB* | 10 | 3 | 97.77% (1492/1526) |
| 1 | *ZFPM2* | 4 | 3 | 90.75% (1491/1643) |
| 1 | *ZFPM2* | 10 | 3 | 97.39% (1493/1533) |
| 2 | - | 0 | 3 | 99.05% (1462/1476) |
| 2 | Mock | 4 | 1 | 100.00% (1475/1475) |
| 2 | Mock | 10 | 2 | 99.19% (1473/1485) |
| 2 | AAVS1 | 4 | 3 | 98.71% (1453/1472) |
| 2 | AAVS1 | 10 | 3 | 98.12% (1461/1489) |
| 2 | *HBB* | 4 | 3 | 98.38% (1454/1478) |
| 2 | *HBB* | 10 | 3 | 97.66% (1458/1493) |
| 2 | *ZFPM2* | 4 | 1 | 100.00% (1474/1474) |
| 2 | *ZFPM2* | 10 | 3 | 98.60% (1476/1497) |
| 3 | - | 0 | 3 | 98.88% (1504/1521) |
| 3 | Mock | 4 | 3 | 98.43% (1504/1528) |
| 3 | Mock | 10 | 3 | 97.54% (1506/1544) |
| 3 | AAVS1 | 4 | 3 | 97.22% (1501/1544) |
| 3 | AAVS1 | 10 | 3 | 97.09% (1501/1546) |
| 3 | *HBB* | 4 | 3 | 98.43% (1504/1528) |
| 3 | *HBB* | 10 | 3 | 97.79% (1503/1537) |
| 3 | *ZFPM2* | 4 | 2 | 98.13% (1522/1551) |
| 3 | *ZFPM2* | 10 | 2 | 99.15% (1519/1532) |

**Supplemental Figure 3:**
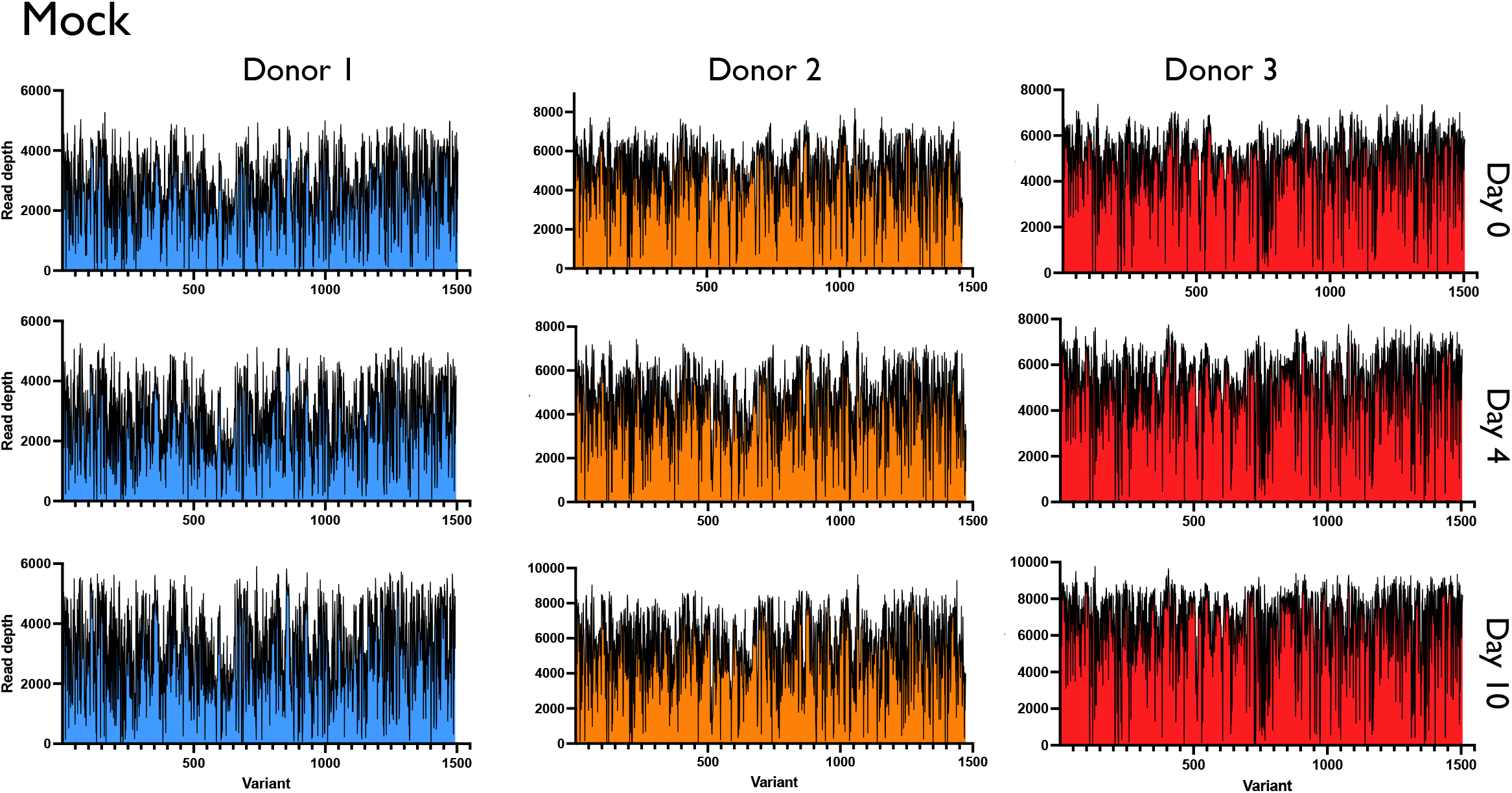
Read depth across variants.

**Supplemental Figure 4:**
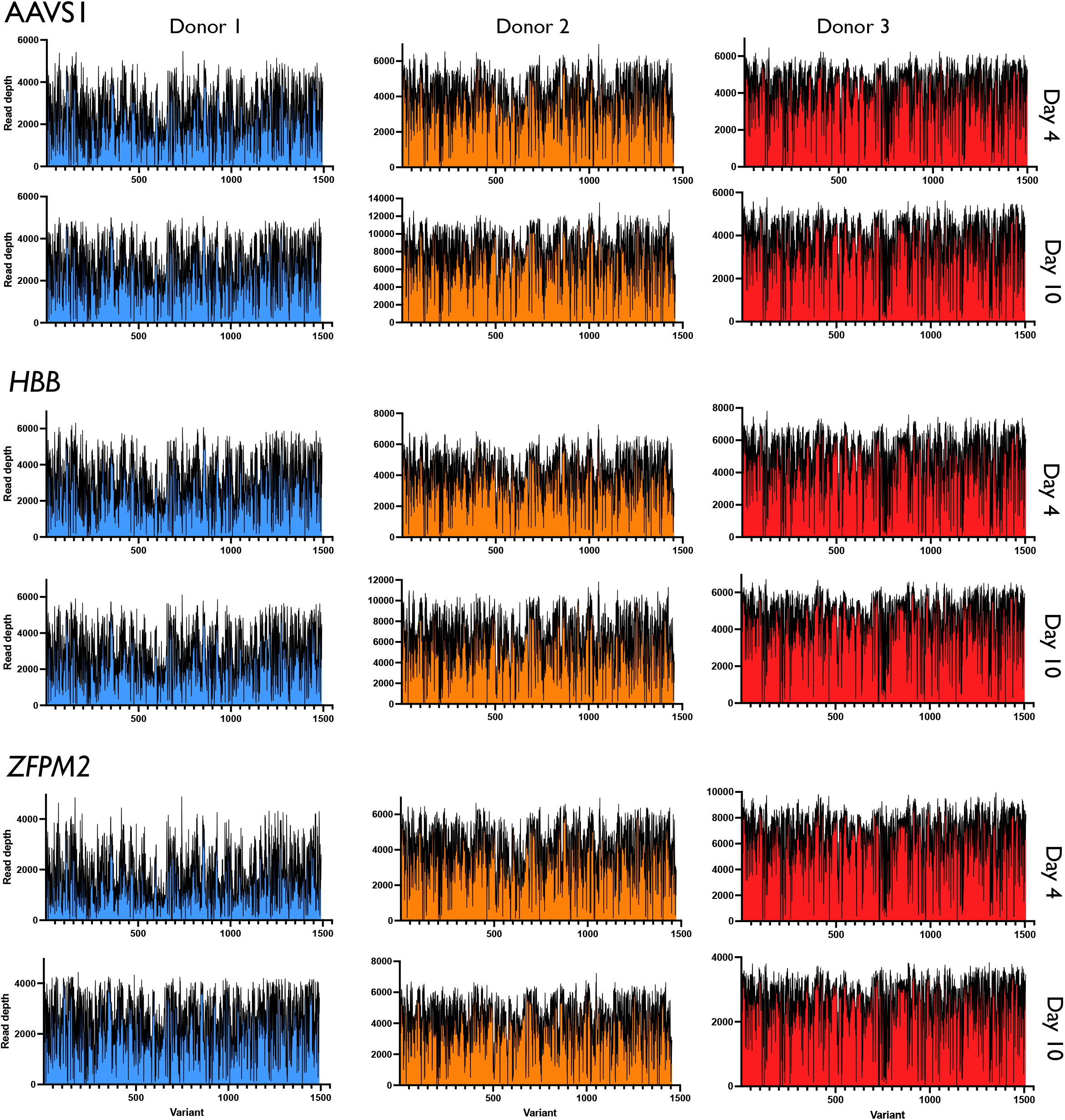
Read depth across variants. Depicted is read depth across all reproducible variants within each treatment and donor. All variants were ordered numerically across genomic coordinates, beginning with chromosome 1 on the left end of the x-axis and ending with the sex chromosomes on the right end of the axis.

**Supplemental Figure 4:**
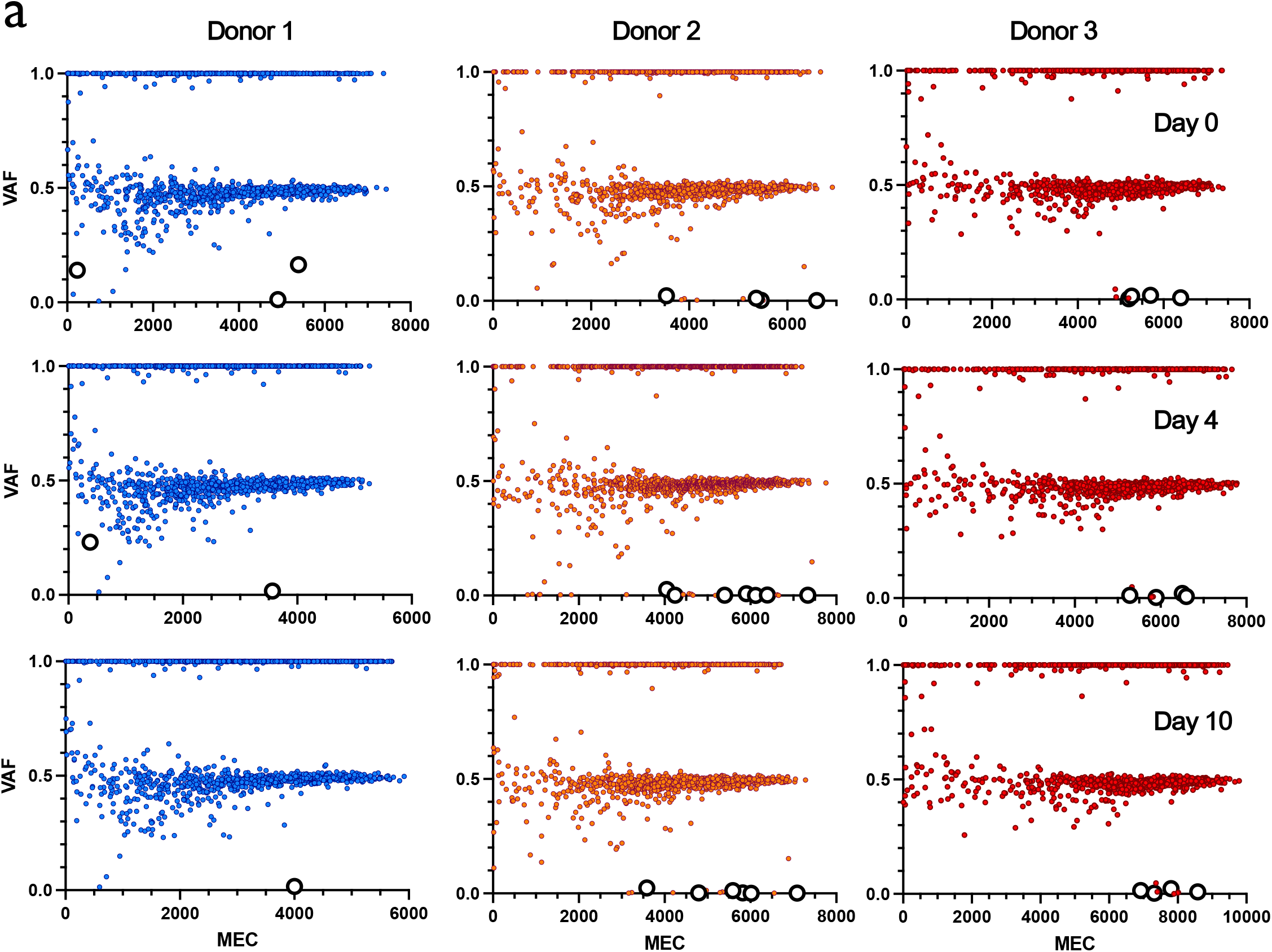
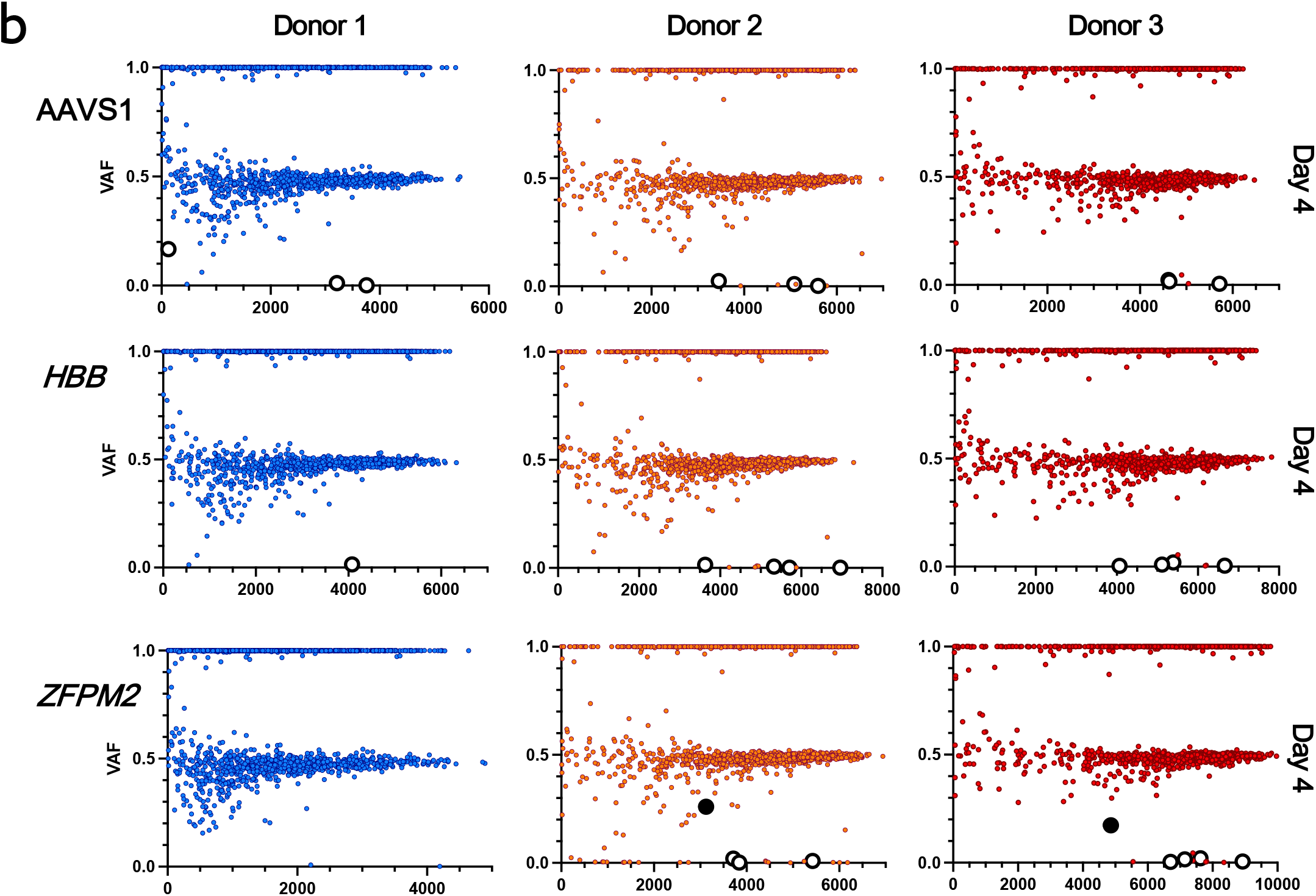
Variants found in treatment groups. a) VAF x MEC for all variants found among technical replicates for Mock treatment for each donor at d0, d4, and d10. Large white points are those that remained after removing germline and synonymous variants. b) VAF x MEC for all variants found among technical replicates for Cas9 treatments for each donor at d4. Large white points are those that remained after removing germline and synonymous variants. Large black points are those that remain after removing variants present in Mock within each donor.

**Supplemental Figure 5:**
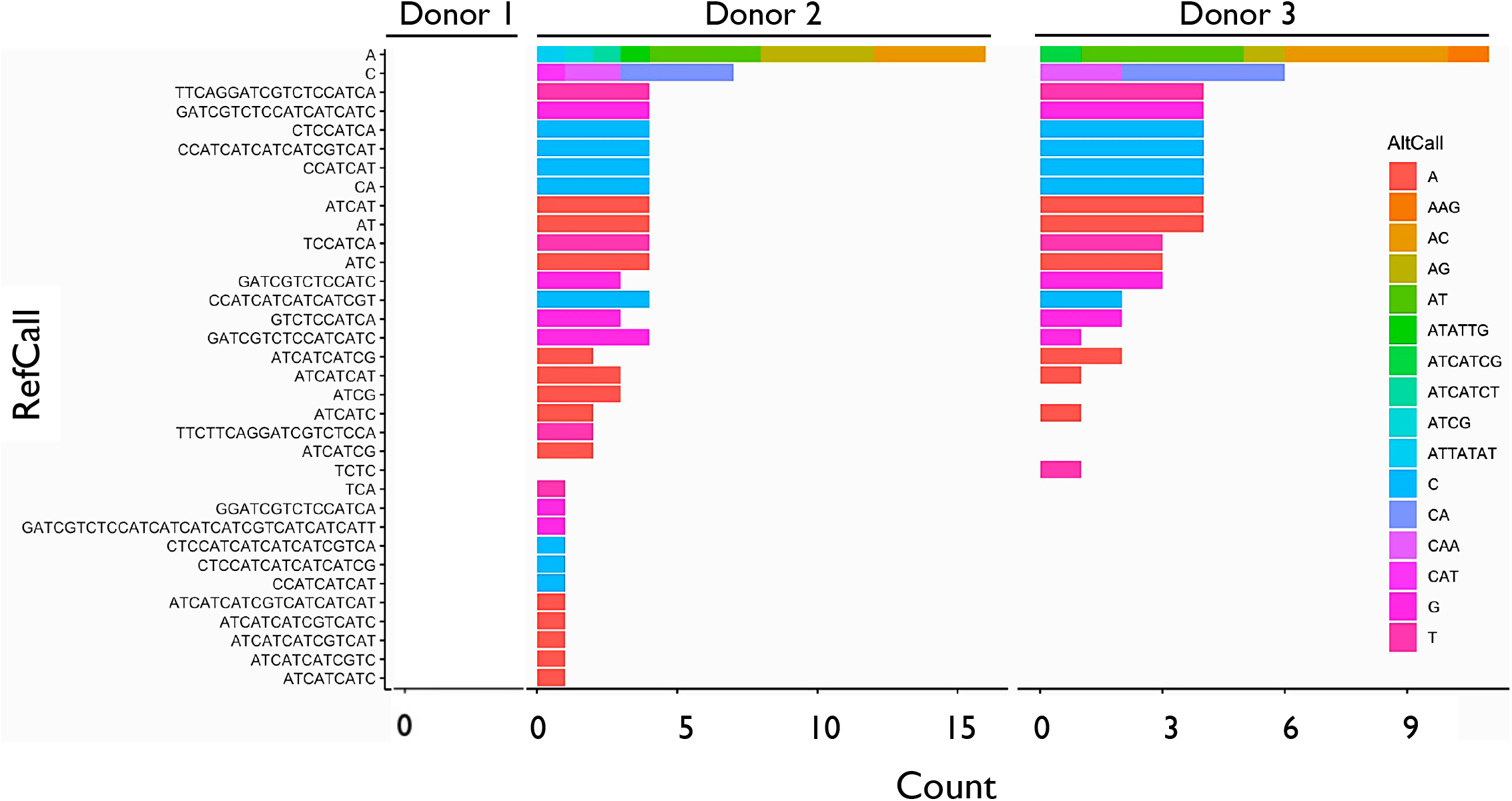
Indel size distribution & frequency for *EZH2*. Specific indels within *EZH2* for each donor in *ZFPM2* treatments (d4 and 10 are combined). Y-axis depicts the normal genome reference call (“RefCall”) and rows depict the frequency at which alternative variant calls (“AltCall”) are found within *EZH2*. Note: no *EZH2* indels detected in Donor 1.

**Supplemental Figure 6:**
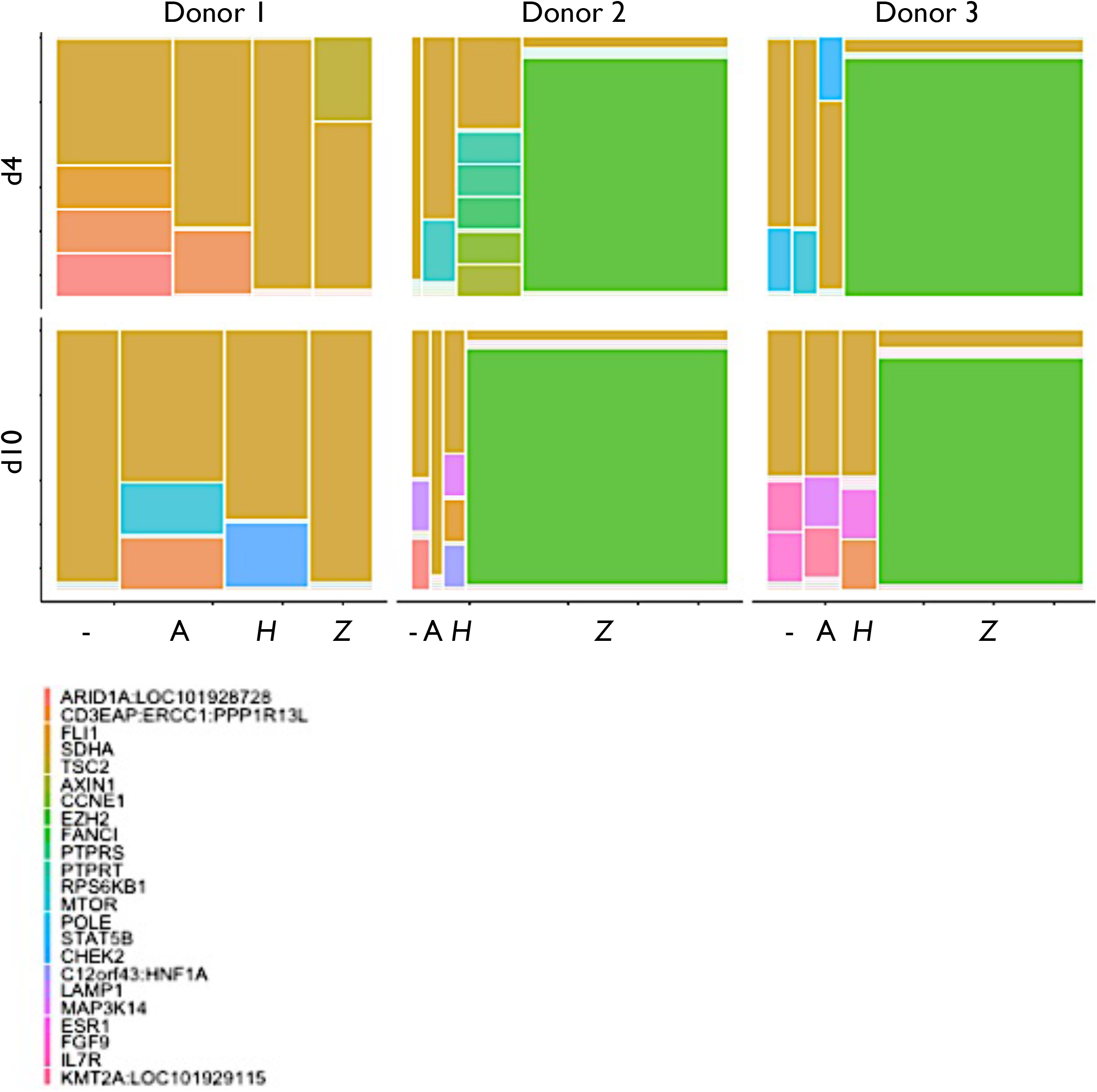
Genes harboring indels among donors, timepoints, & Cas9 treatments. Mosaic plot of genes harboring indels within each donor and Cas9 treatment at d4 and 10 (Mock = −, AAVS1 = A, *HBB* = *H*, and *ZFPM2* = *Z*). Area is proportional to the number of times variants were called within a particular gene within a particular treatment group. Filtering removed germline and synonymous variants as well as MNVs and SNVs.

